# AI-Driven Multiscale Simulations Illuminate Mechanisms of SARS-CoV-2 Spike Dynamics

**DOI:** 10.1101/2020.11.19.390187

**Authors:** Lorenzo Casalino, Abigail Dommer, Zied Gaieb, Emilia P. Barros, Terra Sztain, Surl-Hee Ahn, Anda Trifan, Alexander Brace, Anthony Bogetti, Heng Ma, Hyungro Lee, Matteo Turilli, Syma Khalid, Lillian Chong, Carlos Simmerling, David J. Hardy, Julio D. C. Maia, James C. Phillips, Thorsten Kurth, Abraham Stern, Lei Huang, John McCalpin, Mahidhar Tatineni, Tom Gibbs, John E. Stone, Shantenu Jha, Arvind Ramanathan, Rommie E. Amaro

## Abstract

We develop a generalizable AI-driven workflow that leverages heterogeneous HPC resources to explore the time-dependent dynamics of molecular systems. We use this workflow to investigate the mechanisms of infectivity of the SARS-CoV-2 spike protein, the main viral infection machinery. Our workflow enables more efficient investigation of spike dynamics in a variety of complex environments, including within a complete SARS-CoV-2 viral envelope simulation, which contains 305 million atoms and shows strong scaling on ORNL Summit using NAMD. We present several novel scientific discoveries, including the elucidation of the spike’s full glycan shield, the role of spike glycans in modulating the infectivity of the virus, and the characterization of the flexible interactions between the spike and the human ACE2 receptor. We also demonstrate how AI can accelerate conformational sampling across different systems and pave the way for the future application of such methods to additional studies in SARS-CoV-2 and other molecular systems.

**ACM Reference Format:** Lorenzo Casalino^1†^, Abigail Dommer^1†^, Zied Gaieb^1†^, Emilia P. Barros^1^, Terra Sztain^1^, Surl-Hee Ahn^1^, Anda Trifan^2,3^, Alexander Brace^2^, Anthony Bogetti^4^, Heng Ma^2^, Hyungro Lee^5^, Matteo Turilli^5^, Syma Khalid^6^, Lillian Chong^4^, Carlos Simmerling^7^, David J. Hardy^3^, Julio D. C. Maia^3^, James C. Phillips^3^, Thorsten Kurth^8^, Abraham Stern^8^, Lei Huang^9^, John McCalpin^9^, Mahidhar Tatineni^10^, Tom Gibbs^8^, John E. Stone^3^, Shantenu Jha^5^, Arvind Ramanathan^2∗^, Rommie E. Amaro^1∗^. 2020. AI-Driven Multiscale Simulations Illuminate Mechanisms of SARS-CoV-2 Spike Dynamics. In *Supercomputing ’20: International Conference for High Performance Computing, Networking, Storage, and Analysis. ACM, New York, NY, USA, 14 pages. https://doi.org/finalDOI*

## 1 JUSTIFICATION

We:

- develop an AI-driven multiscale simulation framework to interrogate SARS-CoV-2 spike dynamics,
- reveal the spike’s full glycan shield and discover that glycans play an active role in infection, and
- achieve new high watermarks for classical MD simulation of viruses (305 million atoms) and the weighted ensemble method (600,000 atoms).

## 2 PERFORMANCE ATTRIBUTES

| Performance Attribute | Our Submission |
| --- | --- |
| Category of achievement | Scalability, Time-to-solution |
| Type of method used | Explicit, Deep Learning |
| Results reported on the basis of | Whole application including I/O |
| Precision reported | Mixed Precision |
| System scale | Measured on full system |
| Measurement mechanism | Hardware performance counters, Application timers, Performance Modeling |

## 3 OVERVIEW OF THE PROBLEM

The SARS-CoV-2 virus is the causative agent of COVID19, a world-wide pandemic that has infected over 35 million people and killed over one million. As such it is the subject of intense scientific investigations. Researchers are interested in understanding the structure and function of the proteins that constitute the virus, as this knowledge aids in the understanding of transmission, infectivity, and potential therapeutics.

A number of experimental methods, including x-ray crystallography, cryoelectron (cryo-EM) microscopy, and cryo-EM tomography are able to inform on the structure of viral proteins and the other (e.g., host cell) proteins with which the virus interacts. Such structural information is vital to our understanding of these molecular machines, however, there are limits to what experiments can tell us.

For example, achieving high resolution structures typically comes at the expense of dynamics: flexible parts of the proteins (e.g., loops) are often not resolved, or frequently not even included in the experimental construct. Glycans, the sugar-like structures that decorate viral surface proteins, are particularly flexible and thus experimental techniques are currently unable to provide detailed views into their structure and function beyond a few basic units. Additionally, these experiments can resolve static snapshots, perhaps catching different states of the protein, but they are unable to elucidate the thermodynamic and kinetic relationships between such states.

In addition to the rich structural datasets, researchers have used a variety of proteomic, glycomic, and other methods to determine detailed information about particular aspects of the virus. In one example, deep sequencing methods have informed on the functional implications of mutations in a key part of the viral spike protein [57]. In others, mass spectrometry approaches have provided information about the particular composition of the glycans at particular sites on the viral protein [54, 69]. These data are each valuable in their own right but exist as disparate islands of knowledge. Thus there is a need to integrate these datasets into cohesive models, such that the fluctuations of the viral particle and its components that cause its infectivity can be understood.

In this work, we used all-atom molecular dynamics (MD) simulations to combine, augment, and extend available experimental datasets in order to interrogate the structure, dynamics, and function of the SARS-CoV-2 spike protein (Fig. 1). The spike protein is considered the main infection machinery of the virus because it is the only glycoprotein on the surface of the virus and it is the molecular machine that interacts with the human host cell receptor, ACE2, at the initial step of infection. We have developed MD simulations of the spike protein at three distinct scales, where each system (and scale) is informative, extensive, and scientifically valuable in its own right (as will be discussed). This includes the construction and simulation of the SARS-CoV-2 viral envelope that contains 305 million atoms, and is thus among one of the largest and most complex biological systems ever simulated (Fig. 1A). We employ both conventional MD as well as the weighted ensemble enhanced sampling approach (which again breaks new ground in terms of applicable system size). We then collectively couple these break-through simulations with artificial intelligence (AI) based methods as part of an integrated workflow that transfers knowledge gained at one scale to ‘drive’ (enhance) sampling at another.

**Figure 1:**
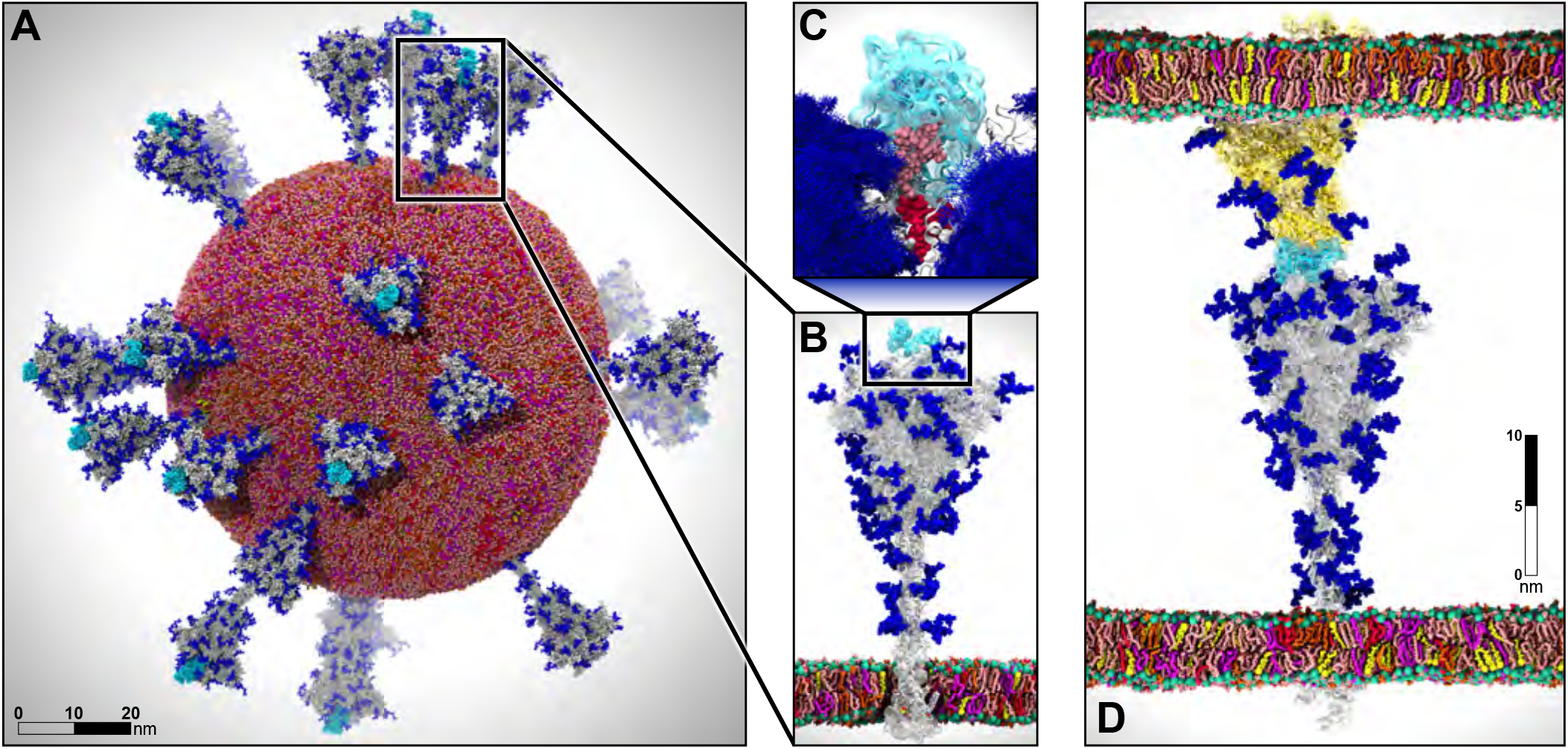
Multiscale modeling of SARS-CoV-2. A) All-atom model of the SARS-CoV-2 viral envelope (305 M atoms), including 24 spike proteins (colored in gray) in both the open (16) and closed states (8). The RBDs in the “up” state are highlighted in cyan) N-/O-Glycans are shown in blue. Water molecules and ions have been omitted for clarity. B) Full-length model of the glycosylated SARS-CoV-2 spike protein (gray surface) embedded into an ERGIC-like lipid bilayer (1.7 M atoms). RBD in the “up” state is highlighted in cyan. C) The glycan shield is shown by overlaying multiple conformations for each glycan collected at subsequent timesteps along the dynamics (blue bushlike representation). Highlighted in pink and red are two N-glycans (linked to N165 and N234, respectively) responsible for the modulation of the RBD dynamics, thus priming the virus for infection. The RBD “up” is depicted with a cyan surface. D) Two-parallel-membrane system of the spike-ACE2 complex (8.5 M atoms). The spike protein, embedded into an ERGIC-like membrane, is depicted with a gray transparent surface, whereas ACE2 is shown with a yellow transparent surface and it is embedded into a lipid bilayer mimicking the composition of mammalian cell membranes. Glycans are shown in blue, whereas water has been omitted for clarity. Visualizations were created in VMD using its custom GPU-accelerated ray tracing engine [23, 58–61].

An additional significant challenge faced in bringing this work to fruition is that it pushes the boundaries of several fields simultaneously, including biology, physics, chemistry, mathematics, and computer science. It is intersectional in nature, and requires the collective work of and effective communication among experts in each of these fields to construct, simulate, and analyze such systems - all while optimizing code performance to accelerate scientific discovery against SARS-CoV-2.

Our work has brought HPC to bear to provide unprecedented detail and atomic-level understanding of virus particles and how they infect human cells. Our efforts shed light on many aspects of the spike dynamics and function that are currently inaccessible with experiment, and have provided a number of experimentally testable hypotheses - some of which have already been experimentally validated. By doing so, we provide new understandings for vaccine and therapeutic development, inform on basic mechanisms of viral infection, push technological and methodological limits for molecular simulation, and bring supercomputing to the forefront in the fight against COVID19.

### 3.1 Methods

#### Full-length, fully-glycosylated spike protein

In this work, we built two full-length glycosylated all-atom models of the SARS-CoV-2 S protein in both closed and open states, fully detailed in Casalino et al [10]. The two all-atom models were built starting from the cryo-EM structures of the spike in the open state (PDB ID: 6VSB [70]), where one receptor binding domain (RBD) is in the “up” conformation, and in the closed state, bearing instead three RBDs in the “down” conformation (PDB ID: 6VXX [66]). Given that the experimental cryo-EM structures were incomplete, the remaining parts, namely (i) the missing loops within the head (residues 16–1141), (ii) the stalk (residues 1141–1234) and (iii) the cytosolic tail (residues 1235–1273), were modelled using MODELLER [51] and I-TASSER [79]. The resulting full-length all-atom constructs were subsequently N-/O-glycosylated using the Glycan Reader & Modeler tool [24] integrated into Glycan Reader [25] in CHARMM-GUI [38]. Importantly, an asymmetric glycoprofile was generated (e.g., not specular across monomers) taking into account the N-/O-glycans heterogeneity as described in the available glycoanalytic data [54, 69]. The two glycosylated systems were embedded into their physiological environment composed of an ERGIC-like lipid bilayer [11, 65] built using CHARMM-GUI [24, 72], explicit TIP3P water molecules [26], and neutralizing chloride and sodium ions at 150 mM concentration, generating two final systems each tallying 1.7 million atoms. Using CHARMM36 all-atom additive force fields [19, 21] and NAMD 2.14 [42], the systems were initially relaxed through a series of minimization, melting (for the membrane), and equilibration cycles. The equilibrated systems were then subjected to multiple replicas of all-atom MD simulation production runs of the open (6x) and closed (3x) systems on the NSF Frontera computing system at the Texas Advanced Computing Center (TACC). A cumulative extensive sampling of ~4.2 and ~1.7 μs was attained for the open and closed systems, respectively. Additionally, a third, mutant system bearing N165A and N234A mutations was built from the open system in order to delete the N-linked glycans and delineate their structural role in the RBD dynamics. This system was also simulated for ~4.2 μs in 6 replicas [10].

#### ACE2-RBD complex MD simulations

The model of the ACE2-RBD complex was based on cryo-EM structure trapping ACE2 as a homodimer co-complexed with two RBDs and B0AT1 transporter (PDB ID 6M17 [73]). Upon removal of B0AT1, ACE2 missing residues at the C terminal end were modeled using I-TASSER [79], whereas those missing at the N terminal end were taken from 6M0J and properly positioned upon alignment of the N terminal helix. Zinc sites including the ions and the coordinating residues were copied from 1R42. The construct was fully N-/O-glycosylated using CHARMM-GUI tools [24, 25, 38] for glycan modeling, reproducing the glycan heterogeneity for ACE2 and RBD reported in the available glycoanalytic data [53, 62, 81]. Similarly, the apo ACE2 homo-dimer was also built upon removal of the RBDs from the holo construct. The glycosylated models were embedded into separate lipid patches with a composition mimicking that of mammalian cellular membranes [11, 65] and simulated in explicit water molecules at 150 mM ion concentration, affording two final systems of ~800,000 atoms each. MD simulations were performed using CHARMM36 all-atom additive force fields [19, 21] along with NAMD 2.14 [42]. The MD protocol was identical to that adopted for the simulation of the full-length spike and it is fully described in Casalino et al [10]. This work is fully detailed in Barros et al [5].

#### Weighted ensemble simulations of spike opening

The spike must undergo a large conformational change for activation and binding to ACE2 receptors, where the receptor binding domain transitions from the “down’,’ or closed state to the “up,” or open state [71]. Such conformational changes occur on biological timescales generally not accessible by classical molecular dynamics simulations [37]. To simulate the full unbiased path at atomic resolution, we used the weighted ensemble (WE) enhanced sampling method [22, 82]. Instead of running one single long simulation, the WE method runs many short simulations in parallel along the chosen reaction coordinates. The trajectories that rarely sample high energy regions are replicated, while the trajectories that frequently sample low energy regions are merged, which makes sampling rare events computationally tractable and gives enhanced sampling. The trajectories also carry probabilities or weights, which are continuously updated, and there is no statistical bias added to the system. Hence, we are able to directly obtain both thermodynamic and kinetic properties from the WE simulations [78].

For this study, the closed model of the glycosylated spike from Casalino et al. [10], was used as the initial structure by only keeping the head domain. The WE simulations were run using the highly scalable WESTPA software [83], with the Amber GPU accelerated molecular dynamics engine [20, 52], version 18. Chamber [13] was used to convert CHARMM36 [19, 21] force fields and parameters from the system developed by Casalino et al. [10] into an Amber readable format. A TIP3P [27] water box with at least 10 Å between protein and box edges was used with 150 mM NaCl, leading the total number of atoms to 548,881. Amber minimization was carried out in two stages. First the solvent was minimized for 10,000 cycles with sugars and proteins restrained with a weight of 100 kcal/mol Å^2^,followed by unrestrained minimization for 100,000 cycles. Next the system was incrementally heated to 300 K over 300 ps. Equilibration and production were carried out in 2 fs timesteps with SHAKE [49] constraints on non-polar hydrogens and NPT ensemble. Pressure and temperature were controlled with Monte Carlo barostat and Langevin thermostat with 1 ps-1 collision frequency. The particle-mesh Ewald (PME) method was used with 10 Å cutoff for non-bonded interactions. The system was first equilibrated for 21 ns of conventional MD. The RMSD of the alpha carbons began to level off around 16 ns, and 24 structures were taken at regular intervals between 16 and 21 ns to use as equally weighted basis states for the WE simulation.

For each WE, tau was set to 100 ps of MD production followed by progress coordinate evaluation, and splitting / merging of walkers and updating weights, with a maximum of 8 walkers per bin. A two dimensional progress coordinate was defined by (*i*): the distance between the center of mass (COM) of the alpha carbons in the structured region of the spike helical core, and the alpha carbons in the four main beta sheets of the RBD (refers to RBD from chain A unless otherwise specified) and (*ii*): the RMSD of the alpha carbons in the four main beta sheets of the RBD to the initial structure (obtained from 1 ns equilibration). This simulation was run for 8.77 days on 80 P100 GPUs on Comet at SDSC collecting a comprehensive sampling of ~7.5 μs, with bin spacing continuously monitored and adjusted to maximize sampling.

After extensive sampling of the RBD closed state, the second progress coordinate was changed to the RMSD of the alpha carbons in the four main beta sheets of the RBD compared to the final open structure, obtained from system 1, after 1 ns of equilibration carried out with identical methods as the closed structure described above, which was initially calculated as 11.5 Å. This allowed more efficient sampling of the transition to the open state by focusing sampling on states which are closer in rotational or translational space to the final state, rather than sampling all conformations that are distinctly different from the closed state. Bin spacing was continuously monitored and adjusted to maximize traversing the RMSD coordinate. The full transition was confirmed when the RMSD coordinate reached below 6 Å and the RBD COM coordinate reached above 8.5 Å (Fig. 2). The simulation was stopped for analysis after 1099 iterations, upon running for 26.74 days on 100 V100 GPUs on Longhorn at TACC and harvesting ~70.0 μs.

**Figure 2:**
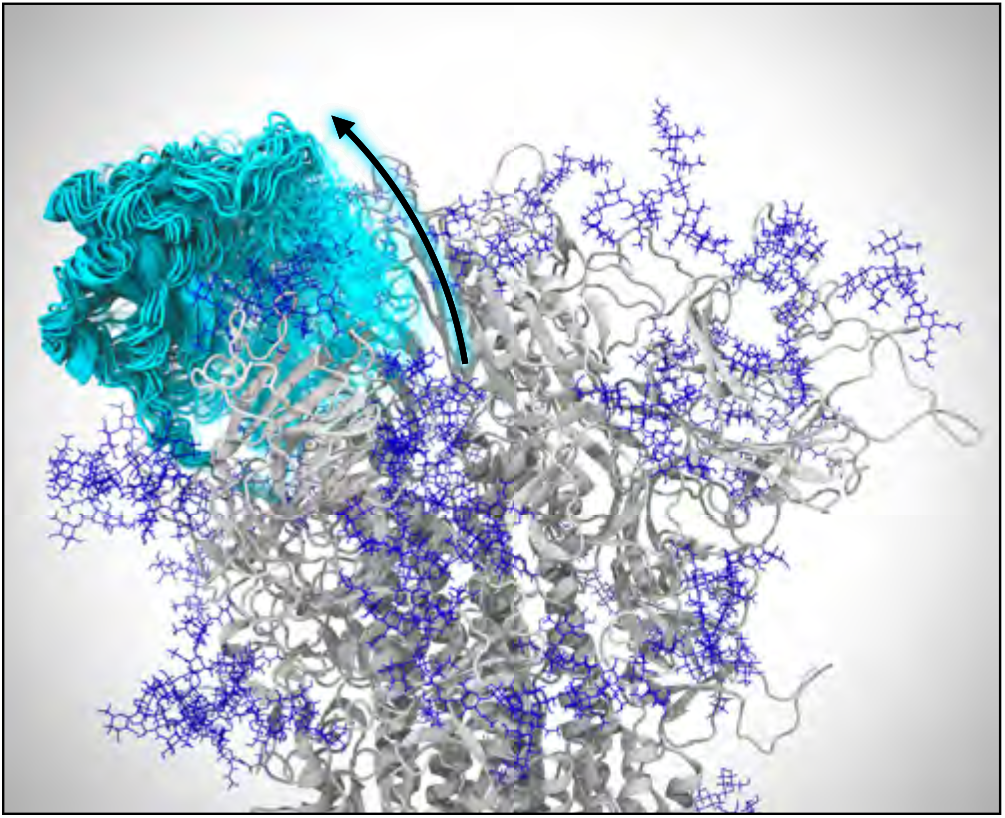
Opening of the spike protein. VMD visualization of weighted ensemble simulations shows the transition of the spike’s RBD from the closed state to the open state. Many conformations of the RBD along its opening pathway are represented at the same time using cyan cartoons and a transparency gradient. Glycans appear as dark blue.

**Figure 3:**
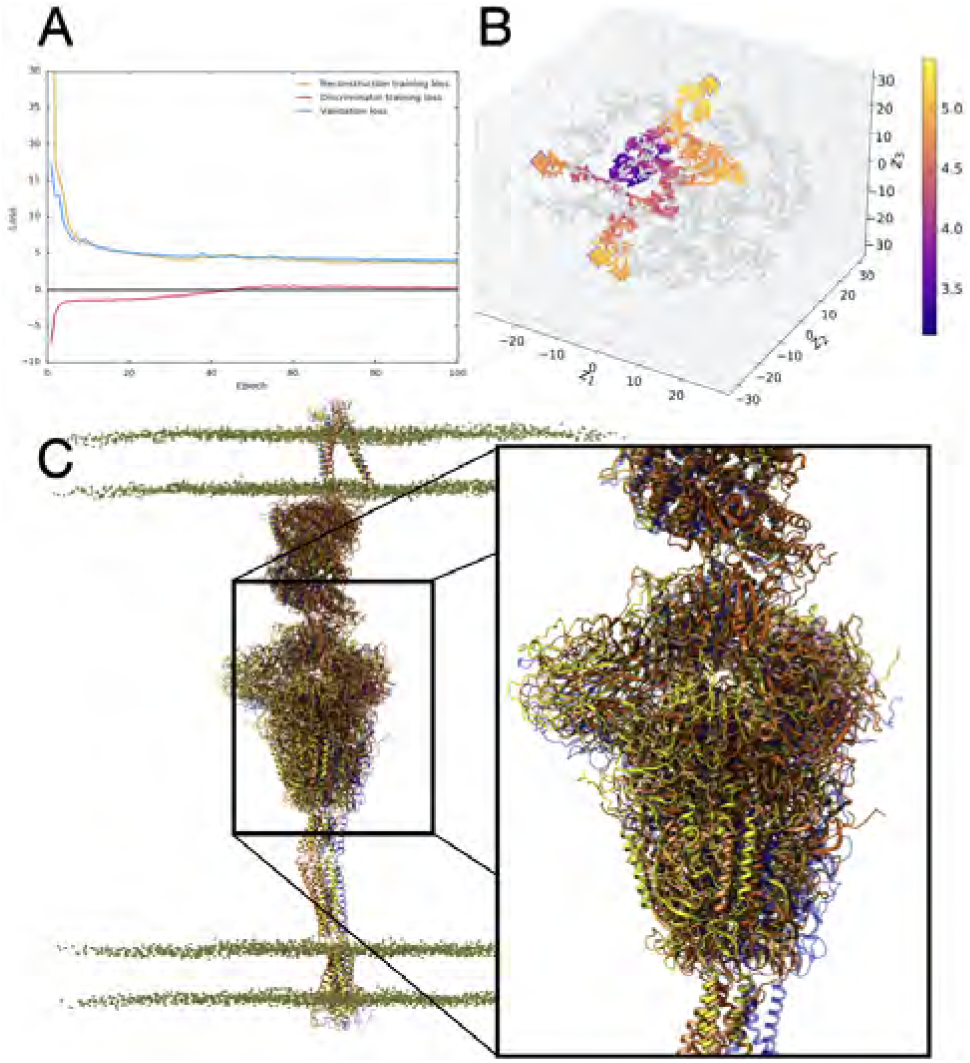
3D-AAE training and test results. A) The loss progression for reconstruction, discriminator and validation loss over 100 epochs. B) The t-SNE plot visualization of the reduced latent space, with training embeddings represented in grey and test examples represented in color over the range of RMSD values. Outliers identified in the outlier detection stage are represented with an outlined diamond. C) VMD visualization of outlier structures (yellow, orange, dark orange) aligned and compared to the starting structure (blue).

A second, independent WE simulation was conducted to determine if the findings of the initial simulation were reproducible, and to use the information on the free energy landscape of the successful transition in the first WE to inform bin spacing and target state definition to run an unsupervised simulation. After 19.64 days on 100 V100 GPUs on TACC Longhorn and ~51.5 μs of comprehensive sampling, successful transitions to the open state were observed, as well as further open states, in which the RBD was observed to be peeling off of the spike core.

#### Two-parallel-membrane system of the spike-ACE2 complex

The SARS-CoV-2 virus gains entry into the host cell through a membrane fusion process taking place upon the recognition of the ACE2 receptors exposed on the host cell. This binding event triggers several, dramatic conformational changes within the spike protein, which becomes primed to pull the two membranes together for fusion, allowing the virus to pour the viral RNA into the host cell. In order to disentangle the mechanistic intricacies underlying this key process, we exploited the wealth of information obtained from the individual simulations described above to assemble an all-atom complex between the full-length spike and the ACE2 dimer. As a first step, equilibrated structures of the spike in the open state and of the ACE2-RBD complex were extracted from their respective individual simulations [5, 9]. Subsequently, the spike protein was superimposed onto the ACE2-RBD complex by aligning the spikes’s RBD “up” with the RBD of the ACE2-RBD complex, allowing for a fairly vertical arrangement of the new construct. In order to preserve the best possible binding interface, the RBD of the spike was discarded, whereas the RBD from the ACE2-RBD complex was retained and linked to the rest of the spike. The spike-ACE2 complex was embedded into a double membrane system: the spike’s transmembrane domain was inserted into a 330 Å × 330 Å ERGIC-like lipid bilayer, whereas for ACE2 a mammalian cellular membrane of the same dimension was used [11, 65]. The two membranes were kept parallel to each other, allowing the use of an orthorhombic box. In order to facilitate the water and ion exchange between the internal and external compartment, an outer-membrane-protein-G (OmpG) porin folded into a beta barrel was embedded into each membrane. The OmpG equilibrated model was obtained from Chen et al [12]. The generated two-membrane construct was solvated with explicit TIP3P water molecules, with the total height of the external water compartment matching the internal one exhibiting a value of 380 Å. Sodium and chloride ions were added at a concentration of 150 mM to neutralize the charge and reshuffled to balance the charge between the two compartments.

The composite system, counting 8,562,698 atoms with an orthorhombic box of 330 Å × 330 Å × 850 Å, was subjected to all-atom MD simulation on the Summit computing system at ORNL using NAMD 2.14 [42] and CHARMM36 all-atom additive force fields [19, 21]. Two cycles of conjugate gradient energy minimization and NPT pre-equilibration were conducted using a 2 fs timestep for a total of ~3 ns. During this phase, the ACE2 and spike proteins and the glycans were harmonically restrained at 5 kcal/mol, allowing for the relaxation of the two lipid bilayers, the OmpG porins, water molecules and ions within the context of the double membrane system. We remark that the two lipid patches were previously equilibrated, therefore not requiring a melting phase at this stage. The dimension of the cell in the xy plane was maintained constant while allowing fluctuation along the z axis. Upon this initial pre-equilibration phase, a ~17 ns NPT equilibration was performed by releasing all the restraints, preparing the system for production run. From this point, three replicas were run or a total of ~522 ns comprehensive simulation time. By using the trained AI learning model, three conformations were extracted from this set of simulations, each of them representing a starting point of a new replica with re-initialized velocities. A total of three additional simulations were therefore performed, collecting ~180 ns and bringing the total simulation time to ~702 ns.

#### SARS-CoV-2 viral envelope

The full-scale viral envelope was constructed using the LipidWrapper program (v1.2) previously developed and described by Durrant et al. [14]. A 350 Å × 350 Å lipid bilayer patch used as the pdb input was generated using CHARMM-GUI with an ERGIC-like lipid composition and an estimated area per lipid of 63 Å. An icospherical mesh with a 42.5 nm radius, in accordance with experimentally-observed CoV-2 radii, was exported as a collada file from Blender (v2.79b) and used as the surface file [31].^1^ LipidWrapper was run in a Python 2.7 conda environment with lipid headgroup parameters “_P,CHL1_O3”, a lipid clash cut-off of 1.0 Å, and filling holes enabled.^2^ The final bilayer pdb was solvated in a 110 nm cubic box using explicit TIP3P water molecules and neutralized with sodium and chloride ions to a concentration of 150 mM. The final system contained 76,134,149 atoms.

Since the LipidWrapper program operates via tessellation, lipid clash removal, and a subsequent lipid patching algorithm, the bilayer output attains a lower surface pressure than that of a bilayer of the same lipid composition at equilibrium [9]. Due to this artifact, as the bilayer equilibrates, the lipids undergo lateral compression resulting in the unwanted formation of pores. Thus, the envelope was subjected to multiple rounds of minimization, heating, equilibration, and patching until the appropriate equilibrium surface pressure was reached.

All-atom MD simulations were performed using NAMD 2.14 and CHARMM36 all-atom additive force fields. The conjugate-gradient energy minimization procedure included two phases in which the lipid headgroups were restrained with 100 and 10 kcal/mol weights, respectively, at 310K for 15,000 cycles each. The membrane was then melted by incremental heating from 25 K to 310 K over 300 ps prior to NPT equilibration. The equilibration sequentially released the harmonic restraints on the lipid headgroups from 100 to 0 kcal/mol over 0.5 ns. Following this sequence, the structure was visually evaluated to determine whether to continue equilibration or to proceed with pore patching. Most structures continued with unrestrained equilibration for 4–26 ns prior to patching, with longer unrestrained equilibrations attributed to later, more stable envelopes.

Patching of the envelope was done by overlapping the initial LipidWrapper bilayer output with the newly-equilibrated envelope. All superimposed lipids within 2.0 Å of the equilibrated lipids were removed to eliminate clashes. Superimposed lipids within 4.0 Å of an equilibrated cholesterol molecule were also removed to eliminate ring penetrations. The patched system, with new lipids occupying the pores, was then re-solvated, neutralized, and subjected to the next round of minimization, heating, and equilibration.

After ten rounds of equilibration and patching, 24 spike proteins with glycans, 8 in the closed and 16 in the open state, were inserted randomly on the envelope using a house tcl script. A random placement algorithm was used in accordance with experimental microscopy imaging which has suggested that there is no obvious clustering of the spikes and no correlation between RBD state and location on the spike surface [31]. The number of spikes was selected based on experimental evidence reporting a concentration of 1000 spikes/nm^2^ on the envelope [31]. The new structure containing spikes was re-solvated, neutralized, and processed to remove clashing lipids prior to further simulation. The resulting cubic solvent box was 146 nm per side and contained 304,780,149 atoms. The spike-inclusive envelope was then subjected to three more equilibration and patching sequences. The final virion used for all-atom MD production runs had a lipid envelope of 75 nm in diameter with a full virion diameter of 120 nm. The complete equilibration of the viral envelope totaled 41 ns on the TACC Frontera system and 75 ns on ORNL Summit. Full-scale viral envelope production simulations were performed on Summit for a total of 84 ns in an NPT ensemble at 310 K, with a PME cutoff of 12 Å for non-bonded interactions.

## 4 CURRENT STATE OF THE ART

### 4.1 Parallel molecular dynamics

NAMD [41] has been developed over more than two decades, with the goal of harnessing parallel computing to create a computational microscope [34, 55] enabling scientists to study the structure and function of large biomolecular complexes relevant to human health. NAMD uses adaptive, asynchronous, message-driven execution based on Charm++[28, 29]. It was one of the first scientific applications to make use of heterogeneous computing with GPUs [43], and it implements a wide variety of advanced features supporting state-of-the-art simulation methodologies. Continuing NAMD and Charm++ developments have brought improved work decomposition and distribution approaches and support for low overhead hardware-specific messaging layers, enabling NAMD to achieve greater scalability on larger parallel systems [32, 44]. NAMD incorporates a collective variables module supporting advanced biasing methods and a variety of in-situ analytical operations [16]. Simulation preparation, visualization, and post-hoc analysis are performed using both interactive and offline parallel VMD jobs [23, 59–61]. NAMD has previously been used to study viruses and large photosynthetic complexes on large capability-oriented and leadership class supercomputing platforms, enabling the high-fidelity determination of the HIV-1 capsid structure [80], the characterization of substrate binding in influenza [15], and the structure and kinetics of light harvesting bacterial organelles [56].

### 4.2 Weighted Ensemble MD simulations

The weighted ensemble (WE) method is an enhanced sampling method for MD simulations that can be orders of magnitude more efficient than standard simulations in generating pathways and rate constants for rare-event processes. WE runs many short simulations in parallel, instead of one long simulation, and directly gives *both* thermodynamic and kinetic properties, which most enhanced sampling methods cannot do. The simulations go through “resampling" where simulations are merged for over-sampled regions and replicated for rare regions so that regions are continuously sampled regardless of energy barriers. The simulations also carry probabilities or “weights" that are continuously updated and no statistical bias is added to the system, so we are able to directly obtain both thermodynamic (e.g., free energy landscape) and kinetic (e.g., rates and pathways) properties from the simulation. In addition, the WE method is one of the few methods that can obtain continuous unbiased pathways between states, so this was the most suitable method for us to obtain and observe the closed to open transition for the spike system. Before the WE method was applied to the spike system under investigation here (about 600,000 atoms), the largest system used for the WE method was the barnase-barnstar complex (100,000 atoms)[50].

### 4.3 AI-driven multiscale MD simulations

A number of approaches, including deep learning methods, have been developed for analysis of long timescale MD simulations [36]. These linear, non-linear, and hybrid ML approaches cluster the simulation data along a small number of latent dimensions to identify conformational transitions between states [6, 46]. Our group developed a deep learning approach, namely the variational autoencoder that uses convolutional filters on contact maps (from MD simulations) to analyze long time-scale simulation datasets and organize them into a small number of conformational states along biophysically relevant reaction coordinates [7]. We have used this approach to characterize protein conformational landscapes [48]. However, with the spike protein, the intrinsic size of the simulation posed a tremendous challenge in scaling our deep learning approaches to elucidate conformational states relevant to its function.

Recently, we extended our approach to adaptively run MD simulation ensembles to fold small proteins. This approach, called DeepDriveMD [35], successively learns which parts of the conformational landscape have been sampled sufficiently and initiates simulations from undersampled regions of the conformational landscape (that also constitute “interesting” features from a structural perspective of the protein). While a number of adaptive sampling techniques exist [2, 8, 30, 33, 47, 67, 68], including based on reinforcement learning methods [39], these techniques have been demonstrated on prototypical systems. In this paper, we utilize the deep learning framework to suggest additional points for sampling and do not necessarily use it in an adaptive manner to run MD simulations (mainly due to the limitations posed by the size of the system). However, extensions to our framework for enabling support of such large-scale systems are straightforward and further work will examine such large-scale simulations.

## 5 INNOVATIONS REALIZED

### 5.1 Parallel molecular dynamics

Significant algorithmic improvements and performance optimizations have been required for NAMD to achieve high performance on the GPU-dense Summit architecture [1, 42, 58]. New CUDA kernels for computing the short-range non-bonded forces were developed that implement a “tile list” algorithm for decomposing the workload into lists of finer grained tiles that more fully and equitably distribute work across the larger SM (streaming multiprocessor) counts in modern NVIDIA GPUs. This new decomposition uses the symmetry in Newton’s Third Law to eliminate redundant calculation without incurring additional warp-level synchronization [58]. CUDA kernels also were added to offload the calculation of the bonded force terms and non-bonded exclusions [1]. Although these terms account for a much smaller percentage of the work per step than that of the short-range non-bonded forces, NAMD performance on Summit benefits from further reduction of CPU workload. NAMD also benefits from the portable high-performance communication layer in Charm++ that communicates using the IBM PAMI (Parallel Active Messaging Interface) library, which improves performance by up to 20% over an MPI-based implementation [1, 32].

Additional improvements have benefited NAMD performance on Frontera. Recent developments in Charm++ now include support for the UCX (Unified Communication X) library which improves performance and scaling for Infiniband-based networks. Following the release of NAMD 2.14, a port of the CUDA tile list algorithm to Intel AVX-512 intrinsics was introduced, providing a 1.8 performance gain over the “Sky Lake” (SKX) builds of NAMD.

A significant innovation in NAMD and VMD has been the development of support for simulation of much larger system sizes, up to two billion atoms. Support for larger systems was developed and tested through all-atom modeling and simulation of the protocell as part of the ORNL CAAR (Center for Accelerated Application Readiness) program that provided early science access to the Summit system [42]. This work has greatly improved the performance and scalability of internal algorithms and data structures of NAMD and VMD to allow modeling of biomolecular systems beyond the previous practical limitation on the order of 250 million atoms. This work has redefined the practical simulation size limits in both NAMD and VMD and their associated file formats, added new analysis methods specifically oriented toward virology [17], and facilitates modeling of cell-scaled billion-atom assemblies, while making smaller modeling projects significantly more performant and streamlined than before [1, 17, 42, 58, 60].

### 5.2 Multiscale molecular dynamics simulations

Often referred to as “computational microscopy,” MD simulations are a powerful class of methods that enable the exploration of complex biological systems, and their time-dependent dynamics, at the atomic level. The systems studied here push state of the art in both their size and complexity. The system containing a full-length, fully-glycosylated spike protein, embedded in a realistic viral membrane (with composition that mimics the endoplasmic reticulum) contains essentially all of the biological complexity known about the SARS-CoV-2 spike protein. The composite system contains ~1.7 million atoms and combines data from multiple cryoEM, glycomics, and lipidomics datasets. The system was simulated with conventional MD out to microseconds in length, and several mutant systems were simulated and validated with independent experiments.

A related set of experiments utilizing the weighted ensemble method, an enhanced sampling technique, explored a truncated version of the spike protein (~ 600,000 atoms with explicit solvent) in order to simulate an unbiased spike protein conformational transition from the closed to open state. This is the largest system, by an order of magnitude, that has been simulated using the WE method (biggest system until now was ~60,000 atoms). Using calculations optimized to efficiently make use of extensive GPU resources, we obtained several full, unbiased paths of the glycosylated spike receptor binding domain activation mechanism.

The second system increases the complexity by an order of magnitude by combining the spike system described above with a full-length, fully-glycosylated model of the ACE2 receptor bound into a host cell plasma membrane. This system represents the encounter complex between the spike and the ACE2 receptor, contains two parallel membranes of differing composition, has both the spike and ACE2 fully glycosylated, and forming a productive binding event at their interface. The composite system contains ~8.5 Million atoms with explicit water molecules and provides unseen views into the critical handshake that must occur between the spike protein and the ACE2 receptor to begin the infection cascade.

Our final system is of the SARS-CoV-2 viral envelope. This system incorporates 24 full-length, fully-glycosylated spike proteins into a viral membrane envelope of realistic (ER-like) composition, where the diameter of the viral membrane is ~80nm and the diameter of the virion, inclusive of spikes, is 146 nm. Until now, the largest system disclosed in a scientific publication was the influenza virus, which contained ~160 million atoms. The SARS-CoV-2 viral envelope simulation developed here contains a composite 305 million atoms, and thus breaks new ground for MD simulations of viruses in terms of particle count, size, and complexity.

Moreover, typical state of the art simulations are run in isolation, presenting each as a self-contained story. While we also do that for each of the systems presented here, we advance on state of the art by using an AI-driven workflow that drives simulation at one scale, with knowledge gained from a disparate scale. In this way, we are able to explore relevant phase space of the spike protein more efficiently and in environments of increasing complexity.

### 5.3 Using AI for driving multiscale simulations

#### Using deep learning to characterize conformational states sampled in the SARS-CoV-2 spike simulations

MD simulations such as the ones described above generate tremendous amounts of data. For e.g., the simulations of the WE sampling of the spike protein’s closed-to-open state generated over 100 terabytes of data. This imposes a heavy burden in terms of understanding the intrinsic latent dimensions along which large-scale conformational transitions can be characterized. A key challenge then is to use the raw simulation datasets (either coordinates, contact matrices, or other data collected as part of a standard MD runs) to cluster conformational states that have been currently sampled, to identify biologically relevant transitions between such states (e.g., open/closed states of spike), and suggest conformational states that may not be fully sampled to characterize these transitions [46].

To deal with the size and complexity of these simulation datasets, approaches that analyze 3D point clouds are more appropriate. Indeed, such approaches are becoming more commonly utilized for characterizing protein binding pockets and protein-ligand interactions. We posited that such representations based on the C^*α*^ representation of protein structures could be viable to characterize large-scale conformational changes within MD simulation trajectories. We leverage the 3D PointNet based [45] adversarial autoencoder (3D-AAE) developed by Zamorski and colleagues [77] to analyze the spike protein trajectories. In this work, we employ the chamfer distance based reconstruction loss and a Wasserstein [4] adversarial loss with gradient penalty [18] to stabilize training. The original PointNet backbone treats the point cloud as unordered, which is true for general point clouds. In our case however, the protein is essentially a 1D embedding into a 3D space. This allows us to define a canonical order of points, i.e. the order in which they appear in the chain of atoms. For that reason, we increase the size-1 1D convolutional encoder kernels from the original PointNet approach to larger kernels up to size 5. This allows the network to not only learn features solely based on distance, but also based on *neighborhood* in terms of position of each atom in the chain. We found that a 4-layer encoder network with kernel sizes [5, 3, 3, 1, 1] and filter sizes [64, 128, 256, 256, 512] performs well for most tasks. A final dense layer maps the vectors into latent space with dimensionality 64. For the generator, we only use unit size kernels with filter dimensions [64, 128, 512, 1024, 3] respectively (the output filter size is always the dimensionality of the problem). The discriminator is a 5 layer fully connected network with layer widths [512, 512, 128, 64, 1].

The trajectories from the WE simulations were used to build a combined data set consisting of 130,880 examples. The point cloud data, representing the coordinates of the 3,375 backbone C^*α*^ atoms of the protein, was randomly split into training (80%) and validation input (20%) and was used to train the 3D-AAE model for 100 epochs using a batch size of 32. The data was projected onto a latent space of 64 dimensions constrained by a gaussian prior distribution with a standard deviation of 0.2. The loss optimization was performed with the Adam optimizer, a variant of stochastic gradient descent, using a learning rate of 0.0001. We also added hyperparameters to scale individual components of the loss. The reconstruction loss was scaled by 0.5 and the gradient penalty by a factor of 10.

The embedding learned from the 3D-AAE model summarizes a latent space that is similar to variational autoencoders, except that 3D-AAEs tend to be more robust to outliers within the simulation data. The embeddings learned from the simulations allow us to cluster the conformations (in an unsupervised manner) based on their similarity in overall structure, which can be typically measured using quantities such as root-mean squared deviations (RMSD).

We trained the model using several combinations of hyperparameters, mainly varying learning rate, batch size and latent dimension. For visualizing and assessing the quality of the model in terms latent space structure, we computed t-SNE [64] dimensionality reductions on the high-dimensional embeddings from the validation set. A good model should generate clusters with respect to relevant biophysical observables not used in the training process. Therefore, we painted the t-SNE plot with the root mean squared deviation (RMSD) of each structure to the starting conformation and observed intelligible clustering of RMSD values. We tested this model on a set of trajectories from the full scale spike-ACE2 system, using the same atom selection (3,375 C^*α*^ atoms) as the corresponding WE spike protein. We subsequently performed outlier detection using the local outlier factor (LOF) algorithm, which uses distance from neighboring points to identify anomalous data. The goal of the outlier detection step is to identify conformations of the protein that are most distinct from the starting structure, in order to story board important events in the transition of the protein from an open to closed conformation. Although the number of outlier conformations detected can be a parameter that the end-user can specify, we selected 20 outlier conformations, based on the extreme LOF scores. These conformations were visualized in VMD [23, 58], and further analyzed using tilt angles of the stalk and the RBD. The final selection included 3 structures which were used as the starting conformations for the next set of simulations. These ‘outlier’ conformers are cycled through additional MD simulations that are driven by the ML-methods.

## 6 HOW PERFORMANCE WAS MEASURED

### 6.1 3D-AAE

Since this application dominantly utilizes the GPU, we do not need to profile CPU FLOPs. Instead, we measure FLOPs for all precisions using the methodology explained in [74] with the NVIDIA NSight Compute 2020 GPU profiling tool. We collect floating point instructions of relevant flavors (i.e. adds, mults, fmas (fused multiply adds) and tensor core operations for FP16, FP32 and FP64) and multiply those with weighting factors of {1, 1, 2, 512} respectively in order to transform those into FLOP counts. The sum of all these values for all precisions will yield our overall mixed precision FLOP count. To exclude FLOPs occuring during initialization and shutdown, we wrap the training iteration loop into start/stop profiler hooks provided by the NVIDIA CuPy Python package.^3^

### 6.2 NAMD

NAMD performance metrics were collected on TACC Frontera, using the Intel msr-tools utilities, with NAMD 2.14 with added Intel AVX-512 support. FLOP counts were measured for each NAMD simulation with runs of two different step counts. The results of the two simulation lengths were subtracted to eliminate NAMD startup operations, yielding an accurate estimate of the marginal FLOPs per step for a continuing simulation [40].

FLOP counts were obtained by reading the hardware performance counters on all CPU cores on all nodes, using the rdmsr utility from msr-tools.^4^ At the beginning of each job, the “TACC stats” system programs the core performance counters to count the 8 sub-events of the Intel FP_ARITH_INST_RETIRED.^5^ Counter values are summed among the 56 cores in each node, and ultimately among each node. Each node-summed counter value is scaled by the nominal SIMD-width of the floating point instruction being counted and the 8 classes are added together to provide the total FLOP count per node. The hardware counters do not take masked SIMD instructions into account. SIMD lanes that are masked-out still contribute to the total FLOPs, however static analysis of the AVX-512-enabled NAMD executable showed that only 3.7% of FMA instructions were masked.

A breakdown of floating point instruction execution frequency for the AVX-512 build of NAMD across 2048 nodes is shown in Table 1. For CPU versions of NAMD, arithmetic is performed in double precision, except for single-precision PME long-range electrostatics calculations and associated FFTs. In the GPU-accelerated NAMD on Summit, single-precision arithmetic is used for both PME and also for short-range non-bonded force calculations, significantly increasing the fraction of single-precision instructions, at the cost of requiring a mixed-precision patch-center-based atomic coordinate representation to maintain full force calculation precision [42, 58].

**Table 1:**
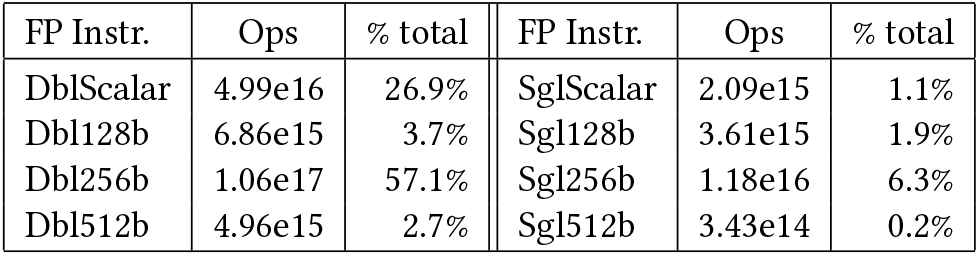
NAMD AVX-512 FP operation breakdown.

## 7 PERFORMANCE RESULTS

### 7.1 3D-AAE training performance

We used the aforementioned recipe for GPU profiling to determine the performance for the 3D-AAE training. We measure the FLOP counts individually for 2 training and 1 validation steps for a batch size of 32. The latent dimension of the model is a free hyperparameter and affects the FLOP count. We trained three models with latent dimensions [32, 64, 128] in order to determine an optimal model for the task and thus we profile and report numbers for all of those. All models were trained for 100 epochs with batch size 32 on a single V100 GPU each. As mentioned above, the train/valdiation dataset split is 80%/20% and we do one validation pass after each training epoch. Thus, we can assume that this fraction translates directly into the FLOP counts for these alternating two stages. Our sustained performance numbers are computed using this weighted FLOP count average and the total run time. In order to determine peak performance, we compute the instantaneous FLOP rate for the fastest batch during training. Note that the 3D-AAE does exclusively use float (FP32) precision. The performance results are summarized in table 2. Although the model is dense linear algebra heavy, it is also rather lightweight so it cannot utilize the full GPU and thus only delivering 20% of theoretical peak performance.

**Table 2:** 3D-AAE training performance on one V100 GPU.

| Latent Dimensions | Peak TFLOP/s | Sustained TFLOP/s |
| --- | --- | --- |
| 32 | 2.96 | 0.97 |
| 64 | 3.16 | 2.28 |
| 128 | 3.13 | 0.91 |

As expected, the peak performance is very consistent between the runs. The big difference in sustained performance between latent dim 64 and the other two models is that the frequency for computing the t-SNE was significantly reduced, i.e. from every epoch to every 5th. The t-SNE computation and plotting happens after each validation in a background thread on the CPU, but the training epochs can be much shorter than the t-SNE time. In that case, the training will stall until the previous t-SNE has completed. Evidently, decreasing the t-SNE frequency reduces that overhead significantly. We expect that the other models would perform similarly if we would have enabled this optimization for those runs as well. The remaining difference in peak vs. sustained performance can be explained by other overhead, e.g. storing embedding vectors, model checkpoints and the initial scaffolding phase. Furthermore, it includes the less FLOP-intensive validation phase whereas the peak estimate is obtained from the FLOP-heavy training phase.

### 7.2 NAMD simulation performance

Low-level NAMD performance measurements were made on the TACC Frontera system, to establish baseline counts of FLOPs per timestep for the four different biomolecular systems simulated as part of this work, summarized in Table 3, with the breakdown of CPU FLOPs described in Table 1. Sustained NAMD performance measurements were obtained using built-in application timers over long production science runs of several hours, including all I/O, and reported in units of nanoseconds per day of simulation time. NAMD sustained simulation performance for the spike-ACE2 complex is summarized for the TACC Frontera and ORNL Summit systems in Table 4 and Fig. 4. NAMD sustained simulation performance, parallel speedup, and scaling efficiency are reported for the full SARS-CoV-2 virion in Table 5. Peak NAMD mixed-precision FLOP rates on ORNL Summit are estimated in Table 6 by combining sustained performance measurements with FLOPs/timestep measurements.

**Table 3:** NAMD simulation floating point ops per timestep.

| NAMD Simulation | Atoms | FLOPS/step |
| --- | --- | --- |
| ACE2-RBD complex | 800k | 21.57 GFLOPS/step |
| Single Spike | 1.7M | 47.96 GFLOPS/step |
| Spike-ACE2 complex | 8.5M | 243.7 GFLOPS/step |
| SARS-CoV-2 virion | 305M | 8.3511 TFLOPS/step |

**Figure 4:**
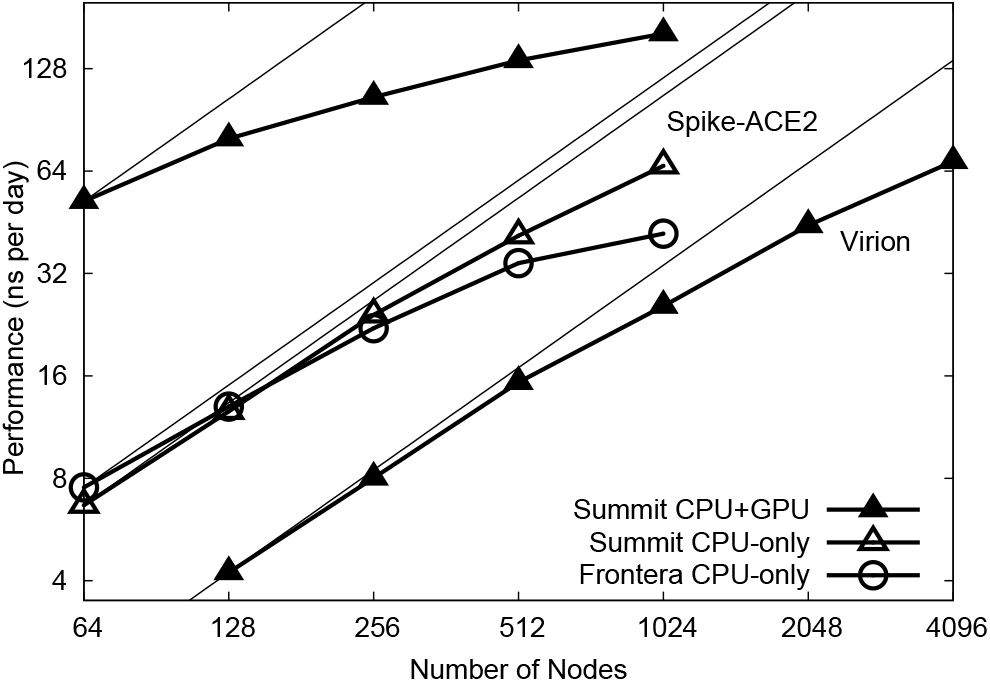
NAMD scaling on Summit and Frontera for 8.5M-atom spike-ACE2 complex (upper lines) and 305M-atom virion (lower line). Thin lines indicate linear scaling.

**Table 4:** NAMD performance: 8.5M-atom Spike-ACE2.

| Nodes | Frontera CPU-only | Summit CPU-only | Summit CPU + GPU |
| --- | --- | --- | --- |
| 64 | 7.52 ns/day | 6.67 ns/day | 52.15 ns/day |
| 128 | 13.00 ns/day | 12.59 ns/day | 79.68 ns/day |
| 256 | 22.09 ns/day | 24.19 ns/day | 105.54 ns/day |
| 512 | 34.32 ns/day | 41.31 ns/day | 135.31 ns/day |
| 1024 | 41.88 ns/day | 66.31 ns/day | 162.22 ns/day |

**Table 5:** NAMD performance: 305M-atom virion.

| Nodes | Summit CPU + GPU | Speedup | Efficiency |
| --- | --- | --- | --- |
| 128 | 4.23 ns/day | ~1.0× | ~100% |
| 256 | 8.02 ns/day | 1.9× | 95% |
| 512 | 15.32 ns/day | 3.6× | 91% |
| 1024 | 25.66 ns/day | 6.1× | 75% |
| 2048 | 44.27 ns/day | 10.5× | 65% |
| 4096 | 68.36 ns/day | 16.2× | 51% |

**Table 6:** Peak NAMD FLOP rates, ORNL Summit

| NAMD Simulation | Atoms | Nodes | Sim rate | Performance |
| --- | --- | --- | --- | --- |
| Spike-ACE2 complex | 8.5M | 1024 | 162 ns/day | 229 TFLOP/s |
| SARS-CoV-2 virion | 305M | 4096 | 68 ns/day | 3.06 PFLOP/s |

## 8 IMPLICATIONS

Our major scientific achievements are:

1. **We characterize for the first time the glycan shield of the full-length SARS-CoV-2 spike protein (including the stalk), and find that two N-glycans linked to N165 and N234 have a functional role in modulating the dynamics of the spike’s RBD.** This unprecedented finding establishes a major new role of glycans in this system as playing an active role in infection, beyond shielding (Fig. 1C) [10].
2. **We discover that the human ACE2 receptor has a flexible hinge in the linker region near the membrane that enables it to undergo exceptionally large angular motions relative to the plane of the membrane.** We predict this flexibility will aid forming productive complexes with the spike protein and may serve as a mechanical energy source during the cell fusion process [5].
3. **We openly share our models, methods, and data, making them freely available to the scientific community.** We are committed to the shared set of principles outlined in Ref. [3]: depositing findings as preprints in advance of formal peer review, making available our models at the time of deposition into a preprint server [5], and releasing the full datasets upon peer review [10]. By doing so, the reproducibility and robustness of our findings and methods are enhanced, and the scientific findings from our simulations are amplified through reuse by others.
4. **We describe for the first time unbiased pathways for the full closed-to-open transition of the spike’s RBD (Fig. 2), where knowledge of this pathway has the potential to inform on mechanisms of viral infection as well as potentially aid in the discovery of novel druggable pockets within the spike.** Our work set a new milestone for the use of the weighted ensemble method in biomolecular simulation, increasing applicable system size by an order or magnitude over current state of the art.
5. **We characterize the spike’s flexibility in the context of ACE2 binding.** One of the most important properties of the spike protein is its intrinsic flexibility, a key feature that facilitates the interaction with the ACE2 receptors exposed on the host cell. Cry-oEM and cryoET structural data revealing the architecture of the SARS-CoV-2 viral particle showed that the spike can tilt up to 60° with respect to the perpendicular to the membrane [31, 75]. Behind this flexibility is the structural organization of the extra-virion portion of the spike, composed of two major domains, the stalk and the head, that are connected through a flexible junction that has been referred to as “hip” (Fig. 5A) [10, 63]. Moreover, the stalk can be further divided into an upper and a lower leg, which correspond to the extra-virion alpha-helices of the coil-coiled trimeric bundle, and the transmembrane domain, which can be intended as the foot of this organizational scaffold. The stalk’s upper leg, lower leg and the foot are interspersed by highly flexible loops defined as “knee” and “ankle” junctions (Fig. 5A) [63]. We then harnessed DeepDriveMD to perform adaptive MD on the Spike-ACE2 8.5 million atoms system. Following this workflow, we extracted three conformations from the first set of Spike-ACE2 MD simulations (replicas 1-3) and subsequently used them as starting points for a new round of MD (replicas 4-6). We then calculated the distribution of the overall spike tilting with respect to the perpendicular to the membrane (Fig. 5E) and of other three angles involving the stalk, namely the “hip” angle between the stalk’s upper leg and the head (Fig. 5B), the “knee” angle between the stalk’s lower and upper legs (Fig. 5C), and the “ankle” angle between the perpendicular to the membrane and the stalk’s lower leg (Fig. 5D). The AI-driven adaptive MD approach expanded the conformational space explored, especially for the knee and hip angles, showing average values of 18.5° ± 7.7° and 13.8° ± 7.6° for replicas 1-3, shifted to 30.4° ± 5.1° and 18.8° ± 6.0° for the subsequent set of MD (replicas 4-6), respectively. The population shift is less pronounced for the ankle, exhibiting an average angle of 21.8° ± 2.7°. These results, in agreement with the data from Turonova et al. [63] that however did not consider the spike in complex with ACE2, reveal large hinge motions throughout the stalk and between the stalk and the head that accommodate the interaction between the spike’s RBD and the ACE2 receptor, preventing the disruption of the binding interface. This is further highlighted by the overall tilting of the spike that remains well defined around 7.3° ± 2.0° (Fig. 5E), showing that the stalk’s inner hinge motions prevent a larger scale bending that could potentially disrupt the RBD-ACE2 interaction.
6. **Our approach points to the very near term ability to accelerate the sampling of dynamical configurations of the complicated viral infection machinery within in the context of its full biological complexity using AI.** The enormous amount of data arising from MD and WE simulations of the single spike served to build and train an AI model using the variational autoencoder deep learning approach, which we demonstrate to accelerate dynamical sampling of the spike in a larger, more complex system (i.e., the two parallel membrane spike-ACE2 complex). Thus, the combination of the AI-driven workflows together with the ground-breaking simulations opens the possibility to overcome a current major bottleneck in the development and use of such ultra-large scale MD simulations, which relates to the efficient and effective sampling of the conformational dynamics of a system with so many degrees of freedom. The scientific implications of such a technological advance, in terms of understanding of the basic science of molecular mechanisms of infection as well as the development of novel therapeutics, are vast.
7. **We establish a new high watermark for the atomic-level simulation of viruses with the simulation of the SARS-CoV-2 viral envelope, tallying 305 million atoms including explicit water molecules, and exhibiting a strong scaling on Summit.** The virion has a realistic ERGIC-like membrane, contains 24 fully glycosylated full-length spikes (in both the open and closed states) and replicates the spatial patterning and density of viral proteins as determined from cryoelectron tomography experiments [31]. These groundbreaking simulations, just now in the process of being fully analyzed, set the stage for future work on SARS-CoV-2 that will be unprecedented in terms of their ability to more closely mimic realistic biological conditions. This includes, for example, the ability to explore the interactions of the virus with multiple receptors on the host cell, or multiple antibodies. It will allow researchers to explore the correlated dynamics of the molecular pieceparts on the surface of the virus and the host cell, and the effects of curvature on such behavior. It will be used as the ground-truth in the development of other simulation approaches, including coarse grained simulation methods, which are under development [76]. It will aid in the development of methods related to the construction of complicated biological membranes [17]. And the list goes on.
8. **We developed an AI-driven workflow as a generalizable framework for multiscale simulation.** Though we focus here on advances made relevant to COVID19, the methods and workflow established here will be broadly applicable to the multiscale simulation of molecular systems.

**Figure 5:**
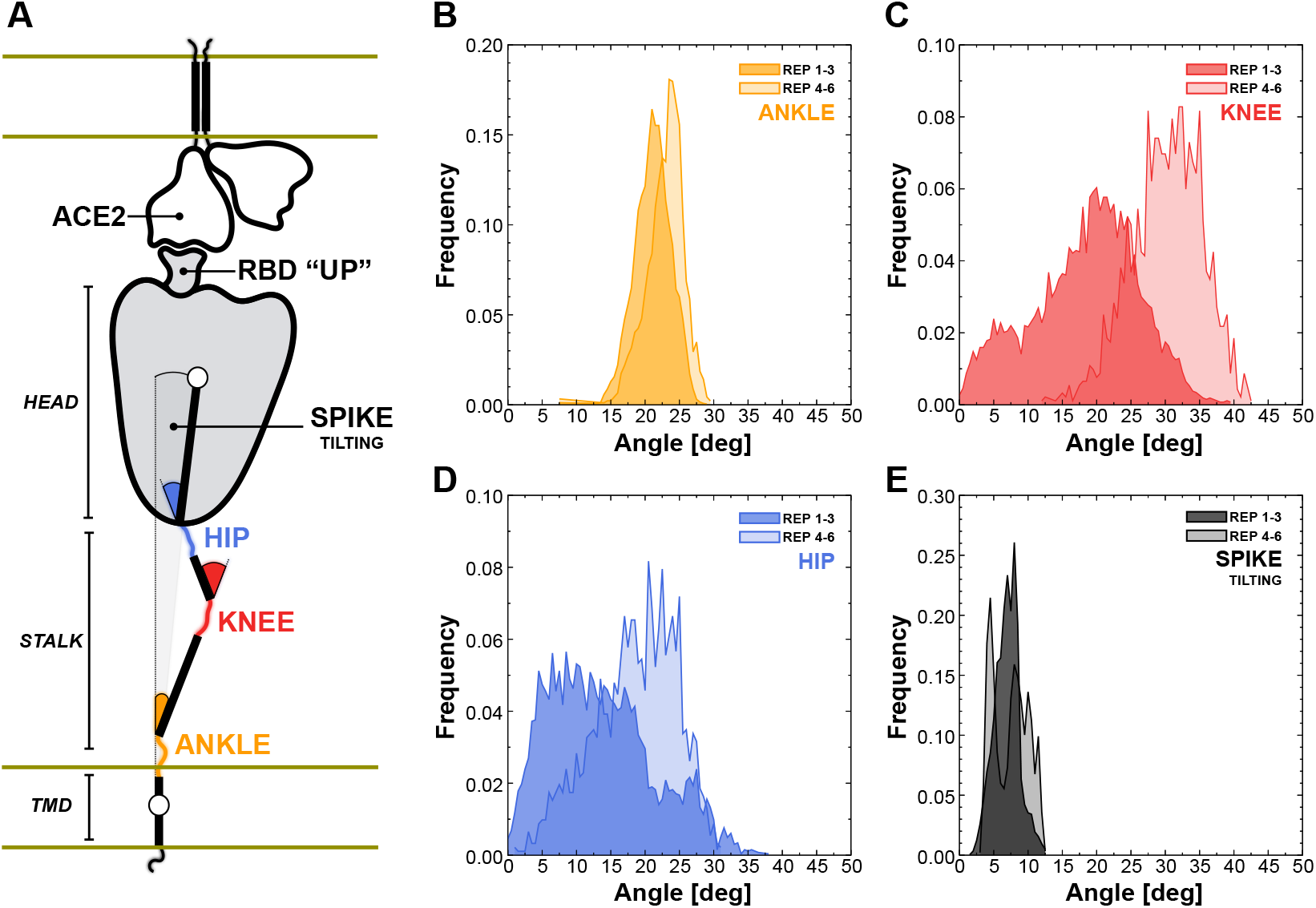
Flexibility of the spike bound to the ACE2 receptor. A) Schematic representation of the two-parallel-membrane system of the spike-ACE2 complex. (B-E) Distributions of the ankle, knee, hip and spike-tilting angles resulting from MD replicas 1-3 (darker color) and 4-6 (lighter color). Starting points for replicas 4-6 have been selected using DeepDriveMD.

## ACKNOWLEDGMENTS

The authors thank D. Maxwell, B. Messer, J. Vermaas, and the Oak Ridge Leadership Computing Facility at Oak Ridge National Laboratory supported by the DOE under Contract DE-AC05-00OR22725. We also thank the Texas Advanced Computing Center Frontera team, especially D. Stanzione and T. Cockerill, and for compute time made available through a Director’s Discretionary Allocation (NSF OAC-1818253). We thank the Argonne Leadership Computing Facility supported by the DOE under DE-AC02-06CH11357. NAMD and VMD are funded by NIH P41-GM104601. The NAMD team thanks Intel and M. Brown for contributing the AVX-512 tile list kernels. Anda Trifan acknowledges support from a DOE CSGF (DE-SC0019323). This work was supported by NIH GM132826, NSF RAPID MCB-2032054, an award from the RCSA Research Corp., a UC San Diego Moore’s Cancer Center 2020 SARS-COV-2 seed grant, to R.E.A. This research was supported by the Exascale Computing Project (17-SC-20-SC), a collaborative effort of the US DOE Office of Science and the National Nuclear Security Administration. Research was supported by the DOE through the National Virtual Biotechnology Laboratory, a consortium of DOE national laboratories focused on response to COVID-19, with funding from the Coronavirus CARES Act. This work used resources, services, and support from the COVID-19 HPC Consortium (https://covid19-hpc-consortium.org/), a private-public effort uniting government, industry, and academic leaders who are volunteering free compute time and resources in support of COVID-19 research. We dedicate this contribution to the memory of Klaus Schulten.

## Footnotes

1 https://www.blender.org/

2 https://docs.anaconda.com/

3 https://cupy.dev/

1 https://github.com/intel/msr-tools

5 https://github.com/TACC/tacc_stats

